# Investigating the interaction between inter-locus and intra-locus sexual conflict using hemiclonal analysis in *Drosophila melanogaster*

**DOI:** 10.1101/2021.10.18.464787

**Authors:** Manas Geeta Arun, Tejinder Singh Chechi, Rakesh Meena, Shradha Dattaraya Bhosle, Srishti, Nagaraj Guru Prasad

## Abstract

Divergence in the evolutionary interests of males and females leads to sexual conflict. Traditionally, sexual conflict has been classified into two types: inter-locus sexual conflict (IeSC) and intra-locus sexual conflict (IaSC). IeSC is modeled as a conflict over outcomes of intersexual reproductive interactions mediated by loci that are sex-limited in their effects. IaSC is thought to be a product of selection acting in opposite directions in males and females on traits with a common underlying genetic basis. While in their canonical formalisms IaSC and IeSC are mutually exclusive, there is growing support for the idea that the two may interact. Empirical evidence for such interactions, however, is limited. Here, we investigated the interaction between IeSC and IaSC in *Drosophila melanogaster*. Using hemiclonal analysis, we sampled 39 hemigenomes from a laboratory-adapted population of *D. melanogaster*. We measured the contribution of each hemigenome to adult male and female fitness at three different intensities of IeSC, obtained by varying the operational sex-ratio. Subsequently, we estimated the intensity of IaSC at each sex-ratio by calculating the intersexual genetic correlation for fitness and the proportion of sexually antagonistic fitness-variation. Our results indicate a statistically non-significant trend suggesting that increasing the strength of IeSC ameliorates IaSC in the population.

## INTRODUCTION

Defined for the first time in 1979 (G. A. Parker, 1979), the term “sexual conflict” is typically used to describe situations which exhibit a negative covariance for fitness between the sexes, i.e., circumstances that are optimal for the fitness of one sex but detrimental to the fitness of the other sex (Schenkel et al., 2018). Examples of sexual conflict encompass a wide range of organisms and traits. They include body size (Stulp et al., 2012), immunocompetence (Sharp & Vincent, 2015; Svensson et al., 2009; Vincent & Sharp, 2014), parental investment (McNamara & Wolf, 2015; Székely, 2014), sex-ratios and sex-allocation (Macke et al., 2014), mating behavior (Culumber et al., 2020), sperm competition (Nandy et al., 2013), traumatic insemination (Dougherty et al., 2017), colour patterns (Price & Burley, 1994), age of maturation (Barson et al., 2015; Mobley et al., 2020) and leaf area (Delph et al., 2011) among others. Conceptually, sexual conflict has been thought to be of two kinds: Interlocus Sexual Conflict (IeSC) or Intralocus Sexual Conflict (IaSC) (Schenkel et al., 2018).

Typically, IeSC has been mathematically modeled as a conflict over mating rates, with male fitness increasing indefinitely with increasing mating rates, while females having an intermediate optimum mating rate (Gavrilets et al., 2001; Rowe et al., 2005). Mating rates are modeled as a function of male and female traits that are sex-limited in their expression (usually called “persistence” and “resistance” traits respectively) (but see Pennell et al. (2016)). Therefore, IeSC is a conflict between a set of loci limited to males, and a different set of loci limited to females. IeSC can also be extended to other spheres of reproductive interactions between males and females; for example, the interplay between the female reproductive tract and male ejaculate components (Sirot et al., 2015). IeSC has been reported in diverse taxa including crickets (Sakaluk et al., 2019), beetles (K. B. McNamara et al., 2020; Wilson & Tomkins, 2014), flatworms (Patlar et al., 2020), snails (Daupagne & Koene, 2020; Swart et al., 2020), and even plants (Lankinen et al., 2016, 2017).

IaSC, on the other hand, is a consequence of males and females sharing the same gene pool while experiencing markedly different selection pressures (Schenkel et al., 2018). IaSC is usually defined for traits that have a common underlying genetic basis in males and females, but have vastly different sex-specific fitness optima (Bonduriansky & Chenoweth, 2009). At the level of a locus, IaSC arises when the allele that is favoured in males is different from the one that is favoured in females (Kidwell et al., 1977). Patterns consistent with IaSC have been reported in a wide range of organisms including guppies (Wright et al., 2018), the bank vole (Lonn et al., 2017), the collared flycatcher (Dutoit et al., 2018), the ant *Nylanderia fulva* (Eyer et al., 2019), and even human beings (Cheng & Kirkpatrick, 2016).

In their traditional formalisms, IaSC (which deals with traits that are shared between the sexes) and IeSC (which deals with traits that are sex-limited in their expression) are mutually exclusive phenomena. However, there have been strong arguments in favour of an interaction between IaSC and IeSC. Pennell & Morrow (2013) argued that IaSC and IeSC could interact in several ways, primarily as a consequence of traits involved in IeSC not being entirely sex-limited in their effects. Traits involved in IeSC could be genetically correlated with traits involved in IaSC. Alternatively, loci involved in IeSC could have pleiotropic effects with negative fitness consequences in the other sex (Pennell et al., 2016). Pennel et al. (2013) also pointed out that processes that resolve IaSC leading to evolution of sexual dimorphism, could trigger IeSC as a result of trait exaggeration. Empirical support for an interaction between IaSC and IeSC is sparse. Working on *Callosobruchus maculates* isofemale lines, Berger et al. (2016) were able to show that multivariate traits associated with high male fitness were genetically associated with a greater drop in line-productivities than could be explained by mate harm (an important aspect of IeSC) or IaSC independently, pointing towards concurrent operation of IaSC and IeSC. However, to the best of our knowledge, no study has yet investigated the consequences of *experimentally* manipulating the intensisty of IeSC on the signal of IaSC in the population. If the assumptions of Pennell et al. (2016) are correct and the loci coding for IeSC traits have negative pleiotropic effects in the opposite sex, then experimentally strengthening IeSC would lead to a strengthening of IaSC in the population.

In the present study, we explored the interaction between IeSC and IaSC in a laboratory adapted population of *Drosophila melanogaster*. *D. melanogaster* is a convenient model organism to address this question as it has been at the forefront of sexual conflict research, primarily because of the tractability of long-term experimental evolution studies using *D. melanogaster*, and the development of crucial genetic tools. One such tool, hemiclonal analysis, which was first developed by Rice, (1996), enables the experimenter to sample hemigenomes from the population of interest and express them in males and females carrying random genetic backgrounds from the population (Abbott & Morrow, 2011). This allows explicit measurements of various quantitative genetic parameters such as additive genetic variances and covariances between quantitative traits, including Darwinian fitness. Using these experimental approaches, *D. melanogaster* has been widely used as a model organism to investigate the evolutionary consequences of IeSC on males and females (Holland & Rice, 1999; Wigby & Chapman, 2004; Nandy et al., 2013a; (Nandy, Gupta, et al., 2013a), quantify genetic variation for IeSC-related traits (Filice & Long, 2016; Linder & Rice, 2005), estimate the intensity of IaSC (Chippindale & Rice, 2001; Collet et al., 2016; Ruzicka et al., 2019), identify traits involved in IaSC (Long & Rice, 2007) and explore sexually antagonistic fitness consequences of male-limited or female-limited evolution (Abbott et al., 2020; Lund-Hansen et al., 2020; Prasad et al., 2007).

To investigate the interaction between IaSC and IeSC, we sampled a panel of hemigenomes from a large laboratory adapted population of *D. melanogaster*. We measured the life-time reproductive fitness of males and females carrying each hemigenome (expressed in a large number of genetic backgrounds randomly sampled from the source population) at three different adult sex-ratios: male-biased (strong IeSC), equal (intermediate IeSC) and female-biased (weak IeSC). Manipulating operational sex-ratios has been one of the two principal techniques of experimentally changing the intensity of IeSC (Janicke & Morrow, 2018; Michalczyk et al., 2011; Nandy, Gupta, et al., 2013a; Nandy et al., 2014; Wigby & Chapman, 2004), the other being experimentally enforcing monogamy (Crudgington et al., 2010; Demont et al., 2014; Gay et al., 2011; Holland & Rice, 1999; Hosken et al., 2001; Tilszer et al., 2006). First, we examined the relationship between the contribution of each hemigenome to sex-specific fitness at each of the three adult sex ratios. Particularly, we attempted to infer if there were any interactions between hemigenome line, sex and sex ratio for fitness. Subsequently, we estimated the following two parameters corresponding to the strength of IaSC for each sex-ratio:

1. Male-female genetic correlation for fitness (r_w,g,mf_): A widely used method of estimating the intensity of IaSC (Bonduriansky & Chenoweth, 2009) with a highly negative r_w,g,mf_ thought to be indicative of strong IaSC (Connallon & Matthews, 2019).
2. The proportion of sexually antagonistic genetic variation (Antagonism Index or AI): A more recent method that partitions fitness-variance along sexually antagonistic and sexually concordant axes (Berger et al., 2014; Ruzicka et al., 2019).

## METHODS

### A. Fly Populations

#### LH

LH is a large laboratory adapted population of *D. melanogaster*. It is a direct descendent of the LHM population used to measure r_w,g,mf_ by Chippindale & Rice (2001), and is related to the populations used by other similar studies (Collet et al., 2016; Ruzicka et al., 2019). The detailed maintenance protocol of LH has been described elsewhere (Nandy et al., 2012). Briefly, it is maintained on a 14 day discrete generation cycle on a standard cornmeal-molasses diet at 25°C, 50% relative humidity, and a 12 hour: 12 hour light-dark cycle. The population consists of a total of 60 vials each containing about 150 eggs in 8-10 ml food. On the 12^th^ day post egg collection, by which time all individuals develop into adult flies, the population is randomly divided into 6 groups of 10 vials each. Flies from each group are mixed together in a flask and subsequently, using light CO_2_ anesthesia, are sorted into 10 food-vials, each containing 16 males and 16 females. Thus, the total population size is 960 females and 960 males spread over 60 vials. Males and females are then allowed to interact for two days in presence of limiting amounts of live yeast. On the 14^th^ day post egg-collection, flies are transferred to fresh food-vials, where they are allowed to lay eggs for 18 hours. The adult flies are then discarded and the eggs are trimmed to a density of 150 per vial. These eggs then start the next generation.

In our experiments, we used the LH population to sample a panel of 39 hemigenomes (see below).

#### LHst

LHst was established by introgressing an autosomal, recessive and benign scarlet eye-colour marker in the LH population. Its maintenance protocol is similar to that of LH, except that the population size is half the population size of LH. LHst is regularly back-crossed to LH to replenish any genetic variation lost due to drift.

#### DxLH

The DxLH population was created by back-crossing the DxIV population (provided to us by Prof. Adam Chippindale) to the LH population for ten generations. DxLH males have a normal X chromosome and a normal Y chromosome. DxLH females have a normal Y chromosome and a compound X chromosome [C(1)DX yf]. This ensures that sons inherit their X chromosome from their father and their Y-chromosome from their mother. Both DxLH males and females have autosomes derived from LH.

#### Clone-generators (CG)

CG males and females have a translocation between the two major autosomes [T(2;3) rdgCst in ripPbwD] (Rice, 1996). CG females have a compound X chromosome [C(1)DX yf] and a Y chromosome. Males have a Y chromosome and an X [snsu(b)] chromosome. CG females enabled us to sample entire haploid genomes (barring the “dot” chromosome 4) and maintain them indefinitely without being damaged by recombination.

### B. Sampling and Maintaining Hemigenomes

We followed a protocol of sampling and maintaining hemigenomes that was similar to the one described by Abbott & Morrow (2011). We chose forty-three males from the LH population randomly. We housed them in separate food-vials with 3 CG females each. From each of the forty-three crosses, we selected one brown-eyed male offspring. Each of these brown-eyed male offspring had a unique haploid “hemigenome” from LH. We then crossed them with 3 CG females each. Absence of molecular recombination in male *D. melanogaster* and the unique features of CG females ensure that the sampled hemigenome gets passed on faithfully from sire to son without being recombined (with the exception of the “dot chromosome”). Each of these 43 crosses represents a unique hemigenome line. We maintained each hemigenome line subsequently by crossing 10 brown-eyed males with 20 CG females every generation. The brown-dominant and scarlet-recessive eye-colour markers on the translocation of the CG females enabled us to distinguish between males that carried the sampled hemigenomes (which were brown-eyed as they were heterozygous for the translocation) and males that were homozygous for the translocation (which were white-eyed). (See Box 2 of Abbott & Morrow (2011) for a detailed schematic.) Four hemigenome lines were lost in an accident. Therefore, we present data from 39 lines.

### C. Fitness Assays

We expressed each hemigenome in males and females carrying the rest of the genome from the LH population and measured their adult fitness at male-biased (8 females: 24 males), equal (16 females: 16 males) and female-biased (24 females: 8 males) sex ratios. Barring the sex-ratios, we tried to ensure that the environment of the fitness assays mimicked the typical LH environment as closely as our experiments could permit.

#### 1. Female Fitness Assay

##### Generating experimental flies

In order to express hemigenomes from each line in females containing a random background from the LH population, we crossed brown-eyed males (heterozygous, carrying the target hemigenome and the translocation) with virgin LH females. To that end, first we collected 30 vials containing 150 eggs each from the LH population. The females emerging from these vials were collected as virgins (within 6 hours of their eclosion) with the help of mild CO_2_ anesthesia by sorting them into vials containing 10 females each. These females were then combined with brown-eyed males from each hemigenome line. For every hemigenome line we set up three to four vials, each containing 5 males from that line and 10 virgin LH females. We allowed these males and females to interact for two days in presence of ad-libitum live yeast (to boost fecundity) and then transferred them to fresh food vials for oviposition for around 18 hours. After discarding the adults, we trimmed the egg-density in each vial to around 250, so that the expected number of larvae surviving in each vial would be around 125 (half the eggs were expected to be unviable). We kept the expected larval density lower than the normal density in the LH population (around 150 per vial) in order to avoid overcrowding in vials that had higher than expected levels of survivorships. Red-eyed females emerging from these vials would be females carrying the target hemigenomes expressed in a random LH background. We refer to these as “focal females”. In order to generate males and competitor females for the assay, we also collected 100 vials of 150 eggs each from the LHst population on the day the eggs from the crosses were trimmed. This ensured that on the day of the experiment all experimental flies were of the same age.

##### Fitness assay

We collected focal females (red-eyed female progeny emerging from the crosses described above) as virgins using light CO_2_ anesthesia and held them in food-vials at a density of 8 females per vial. On the 12^th^ day post egg collection we set up adult competition food-vials supplemented by 100 μL of live-yeast suspension in water. The concentration of the yeast suspension was adjusted according to the sex-ratio treatment such that the per-female yeast availability in the vial was always 0.47 mg. In these adult competition vials, we combined the focal females with competitor LHst females and LHst males at appropriate numbers depending on the sex-ratio treatment. Regardless of the sex-ratio treatment, the total number of flies (males + females) in a vial was always 32, and the ratio of focal females to competitor females was always 1:3. For the male-biased sex-ratio, each vial had 24 LHst males, 2 focal females and 6 LHst competitor females. The equal sex ratio had 16 LHst males, 4 focal females and 12 LHst competitor females in each vial. The female biased sex-ratio had 8 LHst males, 6 focal females and 18 LHst competitor females. We allowed males and females to interact in the adult competition vials for two days. Subsequently, from each vial (regardless of the sex-ratio) we transferred two focal females to a fresh food-vial for egg-laying. We discarded these females after 18 hours and counted the eggs laid in that period, which was used as a measure of the fitness of the focal females in that vial. We performed two replicate assays for each of the sex-ratios, all on separate days. For each replicate assay of each sex-ratio we set up 7 adult competition vials for every hemigenome family. However, due to experimental contingencies, in some cases we had to set up fewer than 7 adult competition vials for some hemigenome lines.

#### 2. Male Fitness Assay

##### Generating experimental flies

The protocol for generating flies for the male fitness assay was similar to the female fitness assay, except that instead of crossing brown-eyed males from each hemigenome line to LH females, we crossed them to virgin DxLH females. This ensured that the red-eyed male progeny emerging from these crosses (the “focal males”) had the target hemigenomes expressed in a random background from the LH population. The eggs laid in the crosses between brown-eyed males from each line and DxLH females were trimmed to a density of around 500 so as to ensure the larval density would be around 125. We also collected 100 vials of 150 eggs each from the LHst population to generate competitor males and females for the fitness assay.

##### Fitness assay

The design of the male fitness assay mirrored that of the female fitness assay. We collected focal males (red-eyed male progeny emerging from the crosses described above) as virgins in food-vials in groups of 8. We also collected as virgins LHst females in groups of 8 per food-vial and competitor LHst males in groups of 6 per vial. On the 12^th^ day post-egg collection we set up adult competition vials as described for the female-fitness experiment. We then combined focal males, competitor LHst males and LHst females in the adult competition vials in appropriate numbers based on the sex-ratio (Male-biased: 6 focal males, 18 LHst competitor males, 8 LHst females; Equal: 4 focal males, 12 LHst competitor males, 16 LHst females; Female-biased: 2 focal males, 6 LHst competitor males, 24 LHst females). We let the flies interact in the adult competition vials for two days. On the 14^th^ day post egg collection, from each vial we transferred 7 randomly chosen LHst females individually into separate test-tubes containing food for oviposition. After 18 hours, we discarded the females and incubated the test tubes in standard maintenance conditions. Twelve days later, when all progeny in the test tubes had developed into adults we froze the test-tubes at −20°C. We scored the progeny from each test-tube for their eye colour. The proportion of red-eyed progeny among all the progeny from the 7 test tubes corresponding to a vial was used as the measure of the fitness of focal males from that vial. For males too, we performed two replicate assays for each of the sex-ratio-treatments, with all six assays being set up separately. Within each assay, for every sex-ratio treatment, we set up 5 adult competition vials for every hemigenome family. In some cases, there were fewer than 5 adult competition vials.

## STATISTICAL ANALYSIS

All analyses were performed in R version 4.0.2.

In order to examine if there was a significant effect of hemigenome line and its interaction with sex and sex ratio, we used the R packages “lme4” and “lmerTest” to fit the following linear mixed effects model on male and female fitness data scaled and centred separately for each day of the experiment:

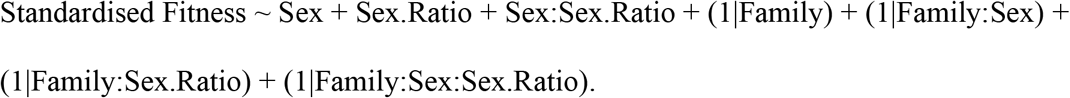

In order to calculate the r_w,g,mf_ we calculated the mean fitness associated with hemigenome line in both males and females. To that end first we arcsin-square-root transformed the male fitness data for each adult competition vial. We divided the data for each day by the mean fitness of that day. Since, we had performed two replicate fitness assays for each sex-ratio with multiple measurements on each day, we calculated the average fitness for hemigenome lines for each sexratio in two steps. For both males and females, for each sex ratio, we first calculated the average fitness for each hemigenome line on each of the two replicate days and then calculated the average of the two averages. We then scaled and centered the data for each sex × sex-ratio combination separately. First, we used this data to calculate genetic correlations for sex-specific fitness across sex ratios. We then calculated the intersexual genetic correlation for fitness (r_w,g,mf_) for each sex-ratio. Following Berger et al. (2014) and Ruzicka et al. (2019), we also calculated the proportion of fitness variation along the sexually antagonistic axis by rotating our original coordinate system represented by a female fitness axis (X-axis) and a male fitness axis (Y-axis) by 45° in the anti-clockwise direction. As a result of this transformation the new X-axis is the axis of sexually concordant fitness variation, while the new Y-axis is the axis of sexually antagonistic fitness variation. We used the following matrix operation separately for the scaled and centered data for each sex-ratio: 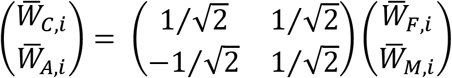, where 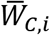 and 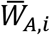 are the sexually concordant and sexual antagonistic fitness components respectively for the hemigenome line i for that sex ratio, and 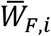 and 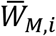 are the average female and male fitnesses respectively for the hemigenome line i for that sex ratio. We then calculated an “antagonism index” (AI) defined by the proportion of variance in fitness lying along the sexually antagonistic axis for each sex ratio.

In order to calculate 95% confidence intervals around our estimates of across sex ratio correlations for sex-specific fitness, r_w,g,mf_ and AI we used a stratified bootstrap approach using the R package “boot” (Canty et al., 2010). For each sex-ratio, we created 10000 data-sets by sampling with replacement within each sex × hemigenome line × day combination. This procedure ensured that each of the bootstrapped data-sets had representation from each sex × hemigenome line × day combination in the same proportions as the original data-set. We also calculated 95% confidence intervals for differences between r_w,g,mf_ and AI estimates of male-biased and female-biased sex ratios to test if they included 0.

Following Ruzicka et al. (2019), we used the R package “MCMCglmm” (Hadfield, 2010) to fit a Bayesian linear mixed effects model using Monte Carlo sampling methods to estimate across sex ratio correlations for sex-specific fitness, r_w,g,mf_ and male and female heritabilities for each sexratio separately. We first scaled and centered arcsin-squareroot transformed male fitness data and female fitness data separately for each day. We fit the following model for each sex-ratio: W_ijkmn_ ~ S_i_ + R_j_ + S_.Rij_ + L_ijk_ + D.L_km_ + ε_ijkmn_, where W_ijkmn_ is the scaled and centered fitness of adultcompetition vial n of sex i, sex ratio j, and hemigenome line k on day m. S_i_, R_j_ and S.R_ij_ represent the fixed effects of sex, sex ratio and their interaction. L_ijk_ represents a term corresponding to the sex-specific random effect of each hemogenome line for sex ratio j. D.L_km_ represents a scalar corresponding to the random interaction of day and hemigenome line. L_ijk_ is modeled to follow a multivariate normal distribution with a mean 0, and whose variance-covariance matrix is given by the additive genetic variance in female fitness (*σ*^2^_*w,g,f*_), and male fitness (*σ*^2^_*w,g,m*_) in each of the three sex ratios; the intersexual genetic covariance for fitness (*Cov_w,g,m,f_*) for each of the three sex ratios; as well as sex-specific genetic covariances for fitness between male biased and female biased sex ratio (*σ*^2^_*w,g,mb–fb*_), between male biased and equal sex ratio (*σ*^2^_w,g,mb–e_), and between female biased and equal sex ratio (*σ*^2^_*w,g,e–fb*_); along with other terms corresponding to genetic covariances for fitness across sex and sex ratios both. ε_ijkmn_ represents the sex and sex-ratio specific residuals. ε_ijkmn_ is modeled to follow a normal distribution with a mean 0 and variance given by the sex and sex-ratio specific residual fitness variance (*σ*^2^_*w,r,m*_ for males and *σ*^2^_*w,r,f*_ for females for each of the three sex-ratios). We used these estimates to calculate the following sex- or sex ratio-specific quantitative genetic parameters:

1. Genetic covariance for fitness between male biased and female biased sex ratio in sex i, 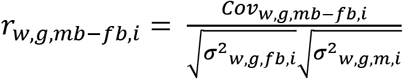
2. Genetic covariance for fitness between male biased and equal sex ratio in sex i, 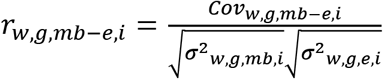
3. Genetic covariance for fitness between equal and female biased sex ratio in sex i, 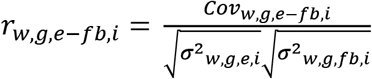
4. Heritability for female fitness in sex ratio j, 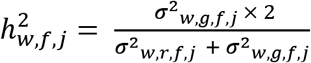
5. Heritability for male fitness in sex ratio j, 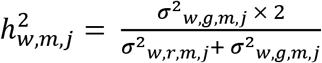
6. Intersexual genetic correlation for fitness in sex ratio j, 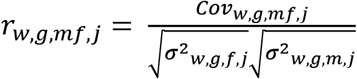

## RESULTS

The output of our linear mixed effects model (Table 1) suggested that there was a significant effect of hemigenome line (likelihood ratio test (LRT), p = 0.0237), its interaction with sex (LRT, p < 0.0001), and the three-way interaction between hemigenome line, sex and sex ratio (LRT, p = 0.0002). While all across-sex ratio correlations for both males and females, and all across-sex correlations for all three sex ratios were positive (Table 2A–2B, Figure 1, Figure 2). many hemigenome lines exhibited fitness rank reversals across sex ratios (Figure 3) or sex (Figure 4), explaining the interactions observed in the linear mixed effects model.

**Figure 1.**
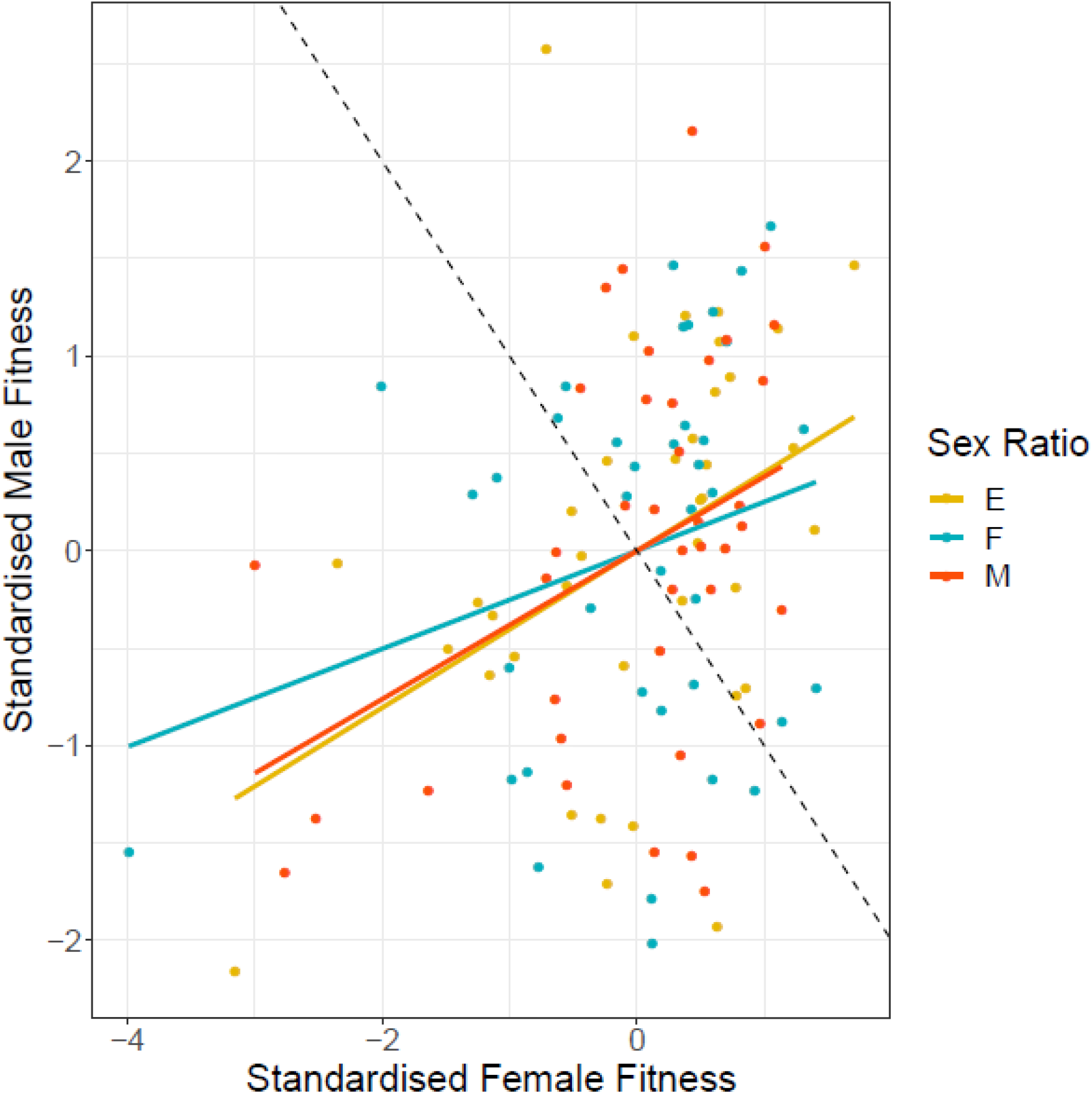
Scaled and centred male and female fitnesses for each of the 39 hemigenome lines for equal sex-ratio (E, yellow), female-biased sex ratio (F, blue) and male-biased sex-ratio (M, red). The solid lines represent the least-squared regression lines for each of the three sexratios. The dashed line represents the axis of sexually antagonistic fitness variation with male-beneficial, female detrimental genptypes to the top-left and female-beneficial, male detrimental genotypees to the bottom-right.

**Figure 2.**
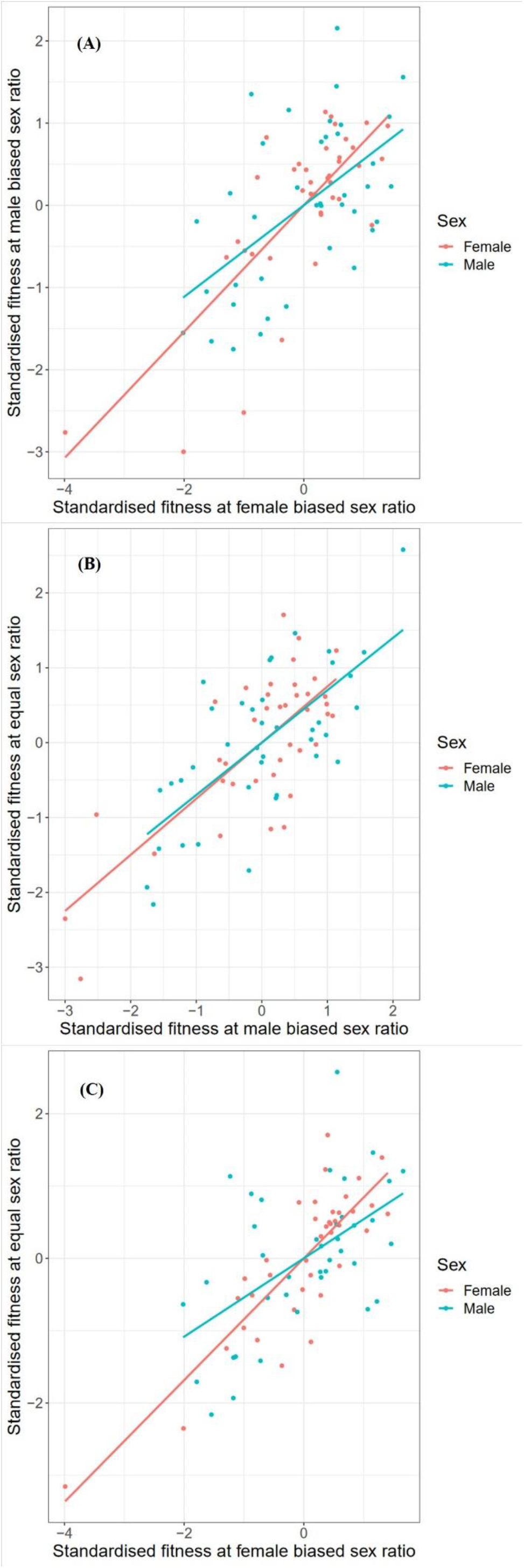
Scatterplots showing standardised male and female fitnesses for various hemigenome lines between (A) male biased and female biased sex ratios, (B) equal and male biased sex ratios, and (C) equal and female biased sex ratios. Blue represents data for males, and red represents data for females. The solid lines represent least-squared regression lines.

**Figure 3.**
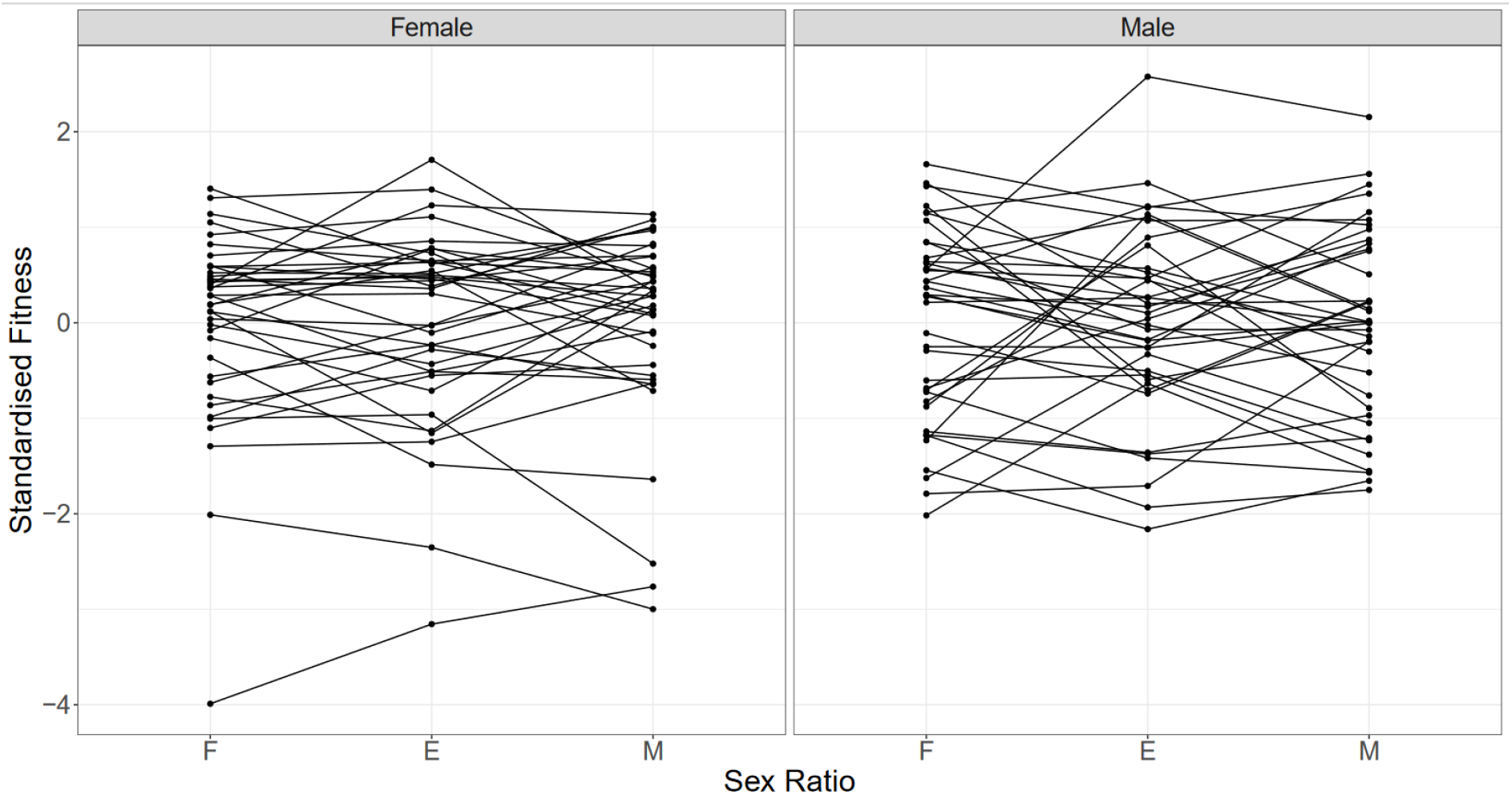
Interaction plots showing standardized fitnesses for various hemigenome lines at female biased (F), equal (E), and male biased (M) sex ratios for females and males. Points connected by a line represent a hemigenome line.

**Figure 4.**
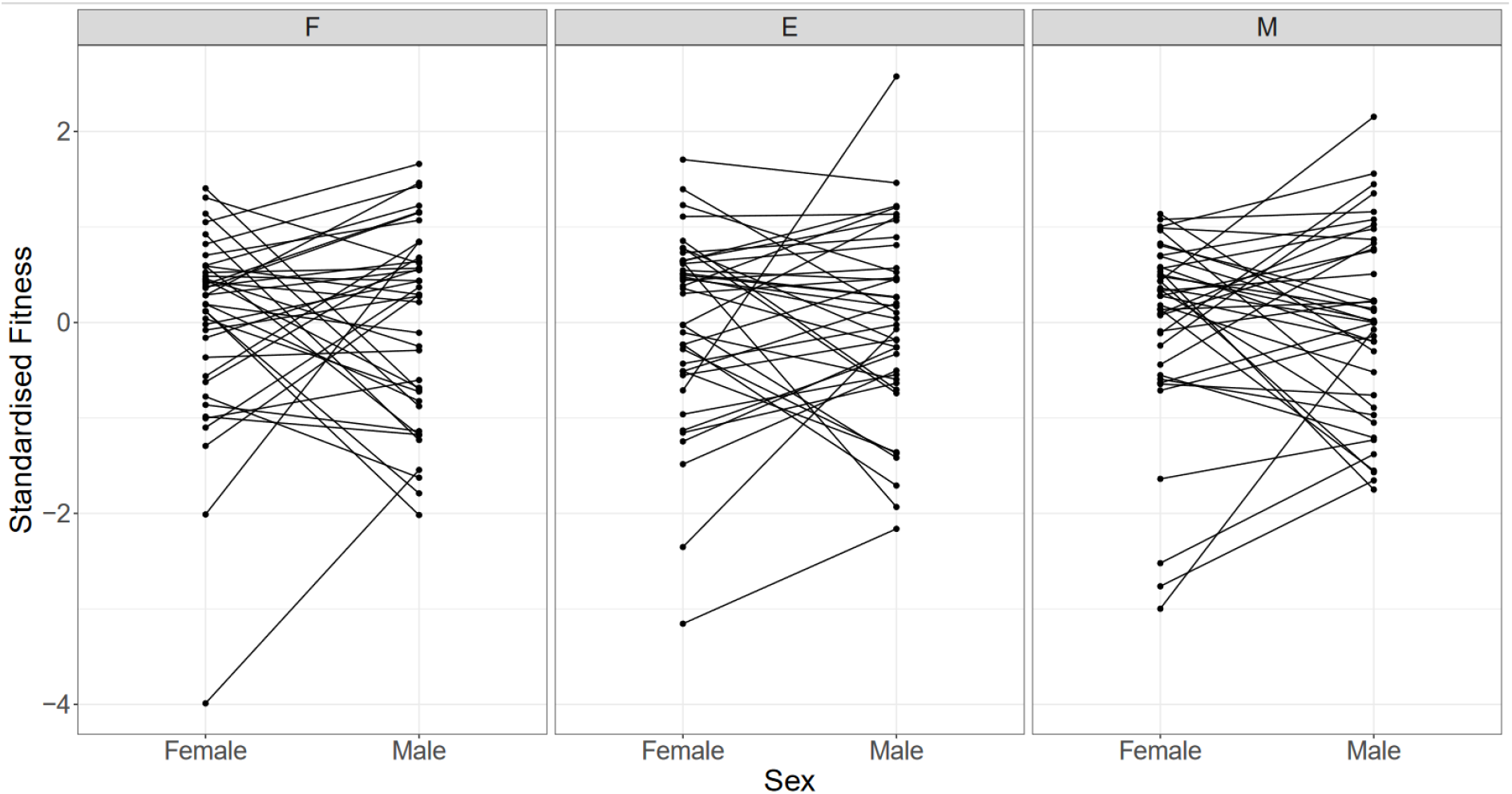
Interaction plots showing standardized fitness for various hemigenome lines for females and males, at female biased (F), equal (E), and male biased (M) sex ratios. Points connected by a line represent a hemigenome line.

**Table 1.**
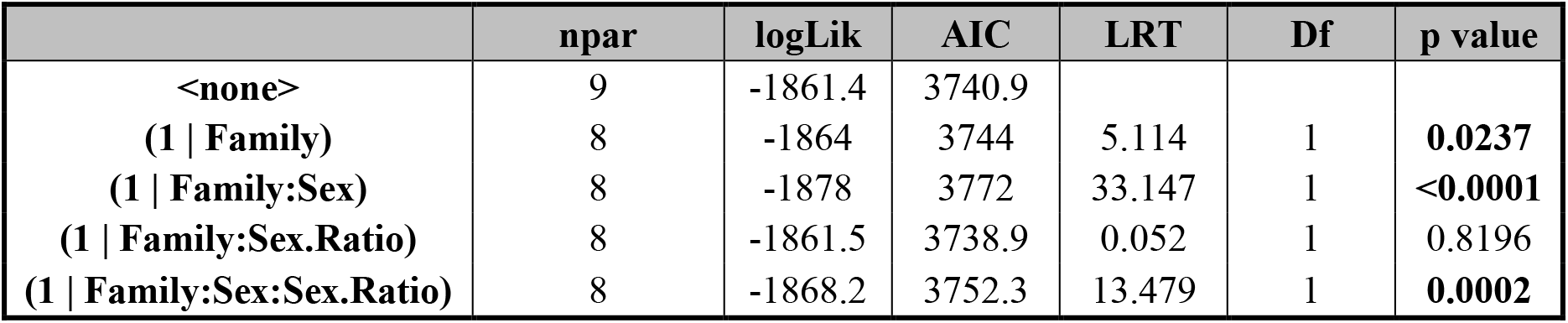
ANOVA-like table for random terms in the linear mixed effects model for male and female fitness

**Table 2.** The summary of results from A) the analysis using hemigenome line averages and B) the MCMCglmm model. Lower and upper CL represent the limits of 95% confidence intervals.

| A) Using Line Averages |  |  |  |  |
| --- | --- | --- | --- | --- |
|  | Sex Ratio | Estimate | Lower CL | Upper CL |
| Intersexual genetic correlation for fitness<br>($r_{w,g,mf}$) | Male Biased | 0.3805 | 0.2992 | 0.5283 |
|  | Equal | 0.4027 | 0.3140 | 0.5526 |
|  | Female Biased | 0.2515 | 0.1198 | 0.4502 |
| Proportion of sexually antagonistic fitness variation (AI) | Male Biased | 0.3097 | 0.2358 | 0.3504 |
|  | Equal | 0.2986 | 0.2237 | 0.3430 |
|  | Female Biased | 0.3742 | 0.2749 | 0.4401 |

|  | Pairs of Sex Ratios | Estimate | Lower CL | Upper CL |
| --- | --- | --- | --- | --- |
| Genetic correlations for female fitness between pairs of sex ratios | Male Biased - Female Biased | 0.7688 | 0.7442 | 0.8497 |
|  | Male Biased - Equal | 0.7493 | 0.7213 | 0.8368 |
|  | Female Biased - Equal | 0.8421 | 0.8403 | 0.8956 |
|  | Pairs of Sex Ratios | Estimate | Lower CL | Upper CL |
| Genetic correlations for male fitness between pairs of sex ratios | Male Biased - Female Biased | 0.5567 | 0.4997 | 0.7262 |
|  | Male Biased - Equal | 0.6995 | 0.6755 | 0.8018 |
|  | Female Biased - Equal | 0.5415 | 0.4664 | 0.7417 |
| B) Using MCMCglmm |  |  |  |  |
|  | Sex Ratio | Estimate | Lower CL | Upper CL |
| Intersexual genetic correlation for fitness<br>($r_{w,g,mf}$) | Male Biased | 0.5056 | 0.1418 | 0.7983 |
|  | Equal | 0.4999 | 0.1397 | 0.7787 |
|  | Female Biased | 0.4462 | 0.0059 | 0.8470 |
| Female Heritability<br>($h_{w,f}^2$) | Male Biased | 0.8702 | 0.5935 | 1.1520 |
|  | Equal | 0.9992 | 0.7337 | 1.2696 |
|  | Female Biased | 0.7385 | 0.5021 | 1.0539 |
| Male Heritability<br>($h_{w,m}^2$) | Male Biased | 0.4788 | 0.2383 | 0.7303 |
|  | Equal | 0.5762 | 0.3192 | 0.8637 |
|  | Female Biased | 0.2229 | 0.0495 | 0.4080 |
|  | Pairs of Sex Ratios | Estimate | Lower CL | Upper CL |
| Genetic correlations for female fitness between pairs of sex ratios | Male Biased - Female Biased | 0.8932 | 0.6888 | 0.9996 |
|  | Male Biased - Equal | 0.8785 | 0.7477 | 0.9994 |
|  | Female Biased - Equal | 0.9536 | 0.8767 | 0.9995 |
|  | Pairs of Sex Ratios | Estimate | Lower CL | Upper CL |
| Genetic correlations for male fitness between pairs of sex ratios | Male Biased - Female Biased | 0.8932 | 0.6888 | 0.9996 |
|  | Male Biased - Equal | 0.9438 | 0.8190 | 1.0000 |
|  | Female Biased - Equal | 0.9010 | 0.7025 | 0.9997 |

The analyses using hemigenome line averages suggested that the r_w,g,mf_ for male-biased sex-ratio (0.3805, 95% CI = [0.2992, 0.52833]) and equal sex ratios (0.4027, 95% CI = [0.3140, 0.5526]) were comparable to each other, but were larger than that for the female-biased sex-ratio (0.2515, 95% CI = [0.1198, 0.4502]). The AI too was comparable for male-biased and equal sex ratios (0.3097, 95% CI = [0.2358, 0.3504] and 0.2986, 95% CI = [0.2237, 0.3430] respectively). The female-biased sex ratio had a higher AI (0.3742, 95% CI = [0.2749, 0.4401]). However, the 95% confidence intervals (CIs) for the difference in the r_w,g,mf_ estimates of male biased and female biased sex ratios (−0.0721, 0.3507) and the 95% CIs for the difference between AI estimates for male biased and female biased sex-ratios (−0.1753, 0.0360) included 0, suggesting these differences were not statistically significant.

The estimates of r_w,g,mf_ from the MCMCglmm model (Table 2B) were slightly higher but the relative trend among sex-ratios was similar. The r_w,g,mf_ estimates were comparable for male-biased (0.5056, 95% credible intervals (CI) = [0.1418, 0.7983]) and equal sex-ratios (0.4999, 95% CI = [0.1397, 0.7787]), while the r_w,g,mf_ estimate for the female-biased sex ratio (0.4462, 95% CI = [0.0059, 0.8470]) was lower (Table 2B). However, the credible interval for the difference between the r_w,g,mf_ estimates for male biased and female biased sex ratios (−0.3788, 0.5561) included 0, suggesting the two were not significantly different. The estimates of female heritabilities in male biased (0.8702, 95% CI = [0.5935, 1.1520]), equal (0.9992, 95% CI = [0.7337, 1.2696]) and female-biased (0.7385, 95% CI = [0.5021, 1.0539]) sex ratios, were higher than the corresponding estimates of male heritabilities at male biased (0.4788, 95% CI = [0.2383, 0.7303]), equal (0.5762, 95% CI = [0.3192, 0.8637]) and female biased (0.2229, 95% CI = [0.0495, 0.4080]) sex ratios. This trend was statistically significant, as the 95% credible intervals for the difference in female and male heritabilities did not overlap with 0 in male biased [-0.7343, −0.0207] and equal [-0.7703, −0.0852] sex ratios, but not in the female biased sex ratio [-0.3740, 0.0668]. Additionally, for both males and females, equal sex-ratio had the highest heritabilities, with the male-biased sex-ratio having marginally lower heritabilities. Both male and female heritabilities were considerably lower in the female-biased sex-ratio. The variance estimate for the interaction between day and hemigenome line was 0.0353 (95% CI = [0.0068, 0.0606]).

## DISCUSSION

We investigated the interaction between inter- and intra-locus sexual conflict in a laboratory adapted population of *D. melanogaster*. We isolated 39 hemigenomes from the LH population and measured the contribution of each hemigenome to the adult lifetime fitness of males and females at male-biased, equal and female-biased sex-ratios. Our analyses yielded the following major findings:

a. At each sex-ratio the intersexual genetic correlation for fitness (r_w,g,mf_) was positive. r_w,g,mf_ was smaller and the antagonism index (AI) higher in the female-biased sex-ratio relative to male-biased or equal sex ratios, suggesting an amelioration of IaSC at higher intensities of IeSC. However, it must be noted that these differences were not statistically significant.
b. Genetic correlations across sex ratios for male and female fitness were strongly positive.
c. There were significant hemigenome line × sex, and hemigenome line × sex × sex ratio interactions for standardized fitness.
d. Heritabilities for fitness were the highest in the equal sex ratio, followed by the male biased sex ratio, and were considerably lower in the female-biased sex-ratio.
e. Estimates of female heritabilities in all three sex ratios were higher than the corresponding estimates of male heritabilities consistent with previous studies on similar systems (Chippindale & Rice, 2001; Innocenti & Morrow, 2010; Collet et al., 2016; Ruzicka et al., 2018).

Below, we discuss some implications of these findings.

Pennell & Morrow (2013) argued that one of the ways in which IaSC and IeSC could interact is if one relaxes the assumption that the loci involved in IeSC are sex-limited in their effects. They pointed out that if traits involved in IeSC have pleiotropic effects in the other sex with negative fitness consequences IaSC could ensue. As a corollary, if one experimentally increases the strength of IeSC in the population, all else being equal, the degree of sexually antagonistic selection (relative to sexually concordant selection) should increase as well. We found no evidence in favour of this prediction. On the contrary, our results suggest a statistically nonsignificant trend in the opposite direction.

Both IaSC and IeSC are complex biological phenomena that involve an interplay of a large number of traits. To be able to predict how changing the intensity of one influences the intensity of the other would, therefore, require an understanding of the genetic correlations between the various traits involved in IaSC and IeSC, and nature of selection acting on each of these traits under different intensities of these conflicts. Below, we describe two plausible scenarios under which strengthening the intensity of IeSC could lead to weaker IaSC within the population.

First, as the intensity of IeSC increases, it is possible that selection gradients on traits involved in IaSC change, leading to a change in the intensity of IaSC over those traits. In an extreme scenario, with increase in the strength of IeSC, one of these selection gradients could change signs in one of the sexes resulting in sexually concordant selection on that trait. Given that we found a strong three-way interaction between sex, sex ratio, and hemigenome line for fitness, in our linear mixed effects model, this explanation becomes fairly plausible. Below, we use available results about locomotory activity to illustrate our point. Adult locomotory activity has been shown to mediate IaSC in *D. melanogaster* (Long & Rice, 2007), with more active males and less active females enjoying higher fitness. Numerous studies have reported patterns that indicate that *D. melanogaster* males that tend to be more active enjoy greater mating success (Hall, 1994; Jordan et al., 2006; Partridge et al., 1987; van Dijken & Scharloo, 1979). On the other hand, female activity stimulates male courtship in *D. melanogaster* (Tompkins et al., 1982). Therefore, active females are thought to attract more courtship from males, resulting in diversion of resources away from egg-production. While a substantial fraction of fitness costs of male-female interactions to females are pre-mating (Partridge & Fowler, 1990), numerous studies have highlighted post-mating fitness costs to females (Fowler & Partridge, 1989; K. Parker et al., 2013; Wigby & Chapman, 2005). Therefore, it is possible that in an environment where IeSC is intense (for example, the male-biased sex-ratio in our experiments), where malecourtship is guaranteed regardless of female activity, selection on females to reduce the number of matings might be stronger than avoiding courtship per se. As a corollary, in an environment with extremely elevated levels of male-courtship, more active females might enjoy higher fitnesses by virtue of their ability to reject male mounting attempts. Therefore, at higher intensities of IeSC, the selection on adult locomotory activity might become sexually concordant reducing the intensity of IaSC. Nandy (2012) evolved replicate populations of *D. melanogaster* at male-biased, equal and female-biased sex-ratios, and reported that both males and females from the male-biased population evolved to become more active than their counterparts evolving under equal and female-biased sex ratios (Nandy et al., 2013b). This suggests that at male-biased sex-ratio, where levels of IeSC are the highest, the IaSC over locomotory activity seems to be weakened, if not removed entirely, so as to permit the evolution of increased locomotory activity levels in both males and females.

Second, increasing the strength of IeSC could ameliorate IaSC if male and female traits (unfortunately called “persistence” and “resistance” traits respectively) involved in IeSC are positively genetically correlated. If the most “resistant” females preferentially mate with the most “persistent” males a positive linkage disequilibrium between “resistance” and “persistence” could build up in the population. As the strength of IeSC increases, by definition, the strength of selection on “persistence” and “resistance” traits increases. If the two sets of traits are positively genetically correlated, this would result in an increase in the strength of sexually concordant selection; all else being equal, this would yield a weakened IaSC signal. Rice et al. (2005) could not find a significant correlation between male and female remating rates in a laboratory population of *D. melanogaster*. However, they did not explicitly observe mating, but measured mating rates in terms of the proportion of females in a vial that remated after their first mating. There are several alternative ways of measuring proxies of persistence and resistance including measuring the latency between the first and the second mating, explicit observations to record matings or measuring courtship related behaviours in males and females. It remains to be explored if these traits are genetically correlated in our panel of hemigenomes.

Our study is also relevant in the context of the “evolutionary inevitability of sexual antagonism” (Connallon & Clark, 2013). Connallon & Clark (2013) used a variant of Fisher’s geometric model to show that as populations adapt to their environments the degree of sexual antagonism in the populations should increase. Consequently, if a population that is well-adapted to its environment is exposed to a novel environment the degree of sexually antagonistic selection experienced by the population should be lower. This idea has been tested in insects by numerous studies with some studies finding evidence in support of the idea (Berger et al., 2014; Long et al., 2012), while others either failed to detect any effect of change of environment on the degree of sexual antagonism (Holman & Jacomb, 2017; Martinossi-Allibert et al., 2018) or reported an increase in sexual antagonism in novel environments (Delcourt et al., 2009; Punzalan et al., 2014). In our case the LH population has been maintained in the laboratory for >500 generations at equal sex-ratio. Therefore, male-biased and female-biased sex-ratios represent novel environments to which the population is not expected to have adapted. Our results are in stark contrast to the idea that maladapted populations should exhibit weaker IaSC. We found that compared to equal sex-ratio, male biased sex ratio exhibited a comparable intensity of IaSC, while the female biased sex-ratio resulted in an *increased* strength of IaSC (lower r_w,g,mf_ and higher IA).

At each of the three sex-ratios our estimates of r_w,g,mf_ were strongly positive. This is in sharp contrast to (Chippindale & Rice, 2001) who had reported a negative r_w,g,mf_ in the ancestral population of the LH population used by us. In fact, several studies have attempted to estimate r_w,g,mf_ in replicates of the original LH_M_ population with different outcomes. Innocenti & Morrow (2010) reported a negative r_w,g,mf_. Collet et al. (2016) compared r_w,g,mf_ across two replicates of the LH_M_ population and reported that one of the replicates had a negative r_w,g,mf_ while the other had an r_w,g,mf_ indistinguishable from 0. Ruzicka et al. (2019) sampled 200 hemogenomes from a replicate of the LHM population and found a positive but non-significant r_w,g,mf_. Ours is the first study to report an r_w,g,mf_ significantly greater than 0. While it is tempting to interpret this as evidence indicating resolution of IaSC through the traditional pathway of sex-specific expression, it might well be a byproduct of strengthening IeSC, as we have argued above. Therefore, further experimental work aimed at understanding the genetic relationships between traits involved in IaSC and IeSC, as well as their selection gradients under various environments is required.

In conclusion, the key findings of our study are as follows:

1. Strengthening the intensity of interlocus sexual conflict did not lead to an increase in the intensity of intralocus sexual conflict, contrary expectations from assumptions in the theoretical literature. We report a non-significant trend in the opposite direction.
2. In contrast with previous studies, we report significantly positive intersexual genetic correlation for fitness, except in the case of female-biased sex ratio, in which case it is indistinguishable from 0 in one of the analyses.
3. Both males and females experience stronger selection in male-biased and equal sex-ratio environments as compared to female-biased sex-ratios.

## Notes

### Competing Interest Statement

The authors have declared no competing interest.

